# A model investigation of short-term synaptic plasticity tuned via Unc13 isoforms

**DOI:** 10.1101/2025.06.18.660326

**Authors:** Magdalena Springer, Stephan Sigrist, Martin Paul Nawrot

## Abstract

Short-term synaptic plasticity (STP) is a fundamental mechanism of neural computation supporting a variety of nervous system functions from sensory adaptation and gain control to working memory and decision making. At the presynaptic release site, an interplay between distinct (M)Unc13 protein isoforms is suggested to orchestrate depressing and facilitating components of STP. In this study, we introduce a modification of the well-established Tsodyks-Markram Model (TMM) for STP. We constrain our model by *in vivo* intracellular recordings in the olfactory system of the fruit fly, where previous work suggested Unc13A to provide a phasic, depressing and Unc13B a tonic, facilitating release component. A combination of a facilitating and a depressing model component indeed allowed for accurate model fits. Differential knock-down experiments of the Unc13A and Unc13B gene variants provide biological model interpretation, linking the protein-specific molecular mechanisms to synaptic function and STP. Our mathematical formulation of protein-dependent STP can be readily and efficiently used to design biologically realistic spiking neural network models that feature different genetically defined synapse types.

---

**S**TP is an ubiquitous phenomenon in excitatory and inhibitory synapses of the invertebrate and vertebrate peripheral and central nervous systems (1–4). It describes the modulation of synaptic efficacy in response to the temporal pattern of presynaptic action potential (AP) firing and takes effect on the time scales of sensory-motor transformation, communication and working memory (3, 5–9). Synaptic computations form fundamental building blocks for network- and systems-level functions. At the synaptic level, STP enables synaptic filtering (10–13), while at the circuit level it supports input-specific sensory adaptation and gain control mechanisms (3, 14–16). Network-level implementations demonstrate its role in shaping attractor dynamics for state transitions (17), maintaining working memory representations (8, 18–20), and facilitating action selection and decision-making processes (8, 20).

New insights into the molecular and cellular mechanisms underlying STP (21, 22) highlight the role of the two protein isoforms Unc13A and Unc13B in *Drosophila melanogaster*. They are enriched at the active zones (AZs) of most chemical synapses and are critical for synaptic vesicle fusion and neurotransmitter release (23). While Unc13A is localized close (≈ 70 nm) to the voltagegated calcium channels by the AZ scaffold protein Bruchpilot, Unc13B is positioned more distantly (≈ 120 nm) by the Syd-1 AZ scaffold protein (24). It has been suggested that Unc13A promotes phasic vesicle release and short-term depression, whereas Unc13B dominated synapses show more tonic vesicle release and dominant short-term facilitation (24–26). The relative expression of the two isoforms varies across different synapse types and thus regulates the short-term dynamics of synaptic transmission and determines the functional synapse type (2, 25, 26).

In mammals, a homologous protein family with distinct Munc13 isoforms has been shown to play a highly similar role in presynaptically regulating synaptic transmission dynamics and STP (2, 27).

## Results

### Application of the TMM to OSN–PN synapses

The TMM (1, 28) is a phenomenological model of STP and finds wide application in local circuit and large scale spiking neural network models (8, 29–31). In its original form (eqns. 1a-1c) it describes a shortterm depressing (STD) component governed by the variable *x* (0 ≤ *x* ≤ 1, *x*_0_ = 1) that represents the fraction of available neurotransmitter resources, and a short-term facilitating (STF) component that is modeled by the variable *u* (0 ≤ *u* ≤ 1, *u*_0_ = 0), representing the AP induced accumulation of calcium in the presynaptic terminal. With each AP of the presynaptic neuron, the facilitation variable *u* increases while the depression variable *x* decreases. The synaptic current *I*_*S*_ is then defined according to

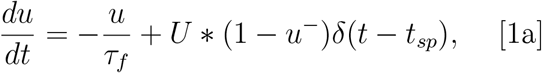

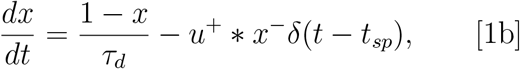

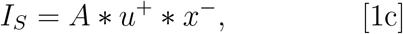

where 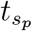 denotes the AP times, *U* is the increment of *u* induced by each AP, *τ*_*d*_ and *τ*_*f*_ represent the time constants with which variables *u* and *x* approach their initial values, and *A* defines the maximum synaptic efficacy. The superscripts − and + indicate the time point right before and right after 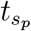, respectively. The parameter set (*A, U, τ*_*d*_, *τ*_*f*_) determines the extend to which facilitation and depression are expressed in the model synapse.

We applied the TMM to electrophysiological data recorded at the synapse between Olfactory Sensory Neurons (OSNs) and Projection Neurons (PNs) in the antennal lobe of the wild-type fruit fly *D. melanogaster* (25). As sketched in Fig. 1 A, whole-cell voltage clamp recordings were performed from individual postsynaptic PNs during electrical stimulation of presynaptic OSN axons in the antennal nerve. The stimulation protocol (Fig. 1 B) included a step change from a regular 10 Hz stimulation to a regular 50 Hz stimulation. The excitatory postsynaptic current (EPSC) amplitude decays over repeated stimulation, indicating a clear depressing component (black curve in Fig. 1 C). At the same time, a high increase in EPSC amplitude after the first short-latency stimulus indicates a component of paired-pulse facilitation at this synapse type. We then fitted the TMM to our experimental data, identifying the best-matching parameter set for each individual neuron (Tab. 1, Material and Methods). The resulting model performance remained poor, despite an extensive parameter search. It particularly could not accurately reproduce the temporal pattern of EPSC amplitudes, failing to simultaneously capture both, the clear depression of high amplitudes and the paired-pulse facilitation in response to the first short inter-stimulus interval (Fig. 1 C).

**Fig. 1.**
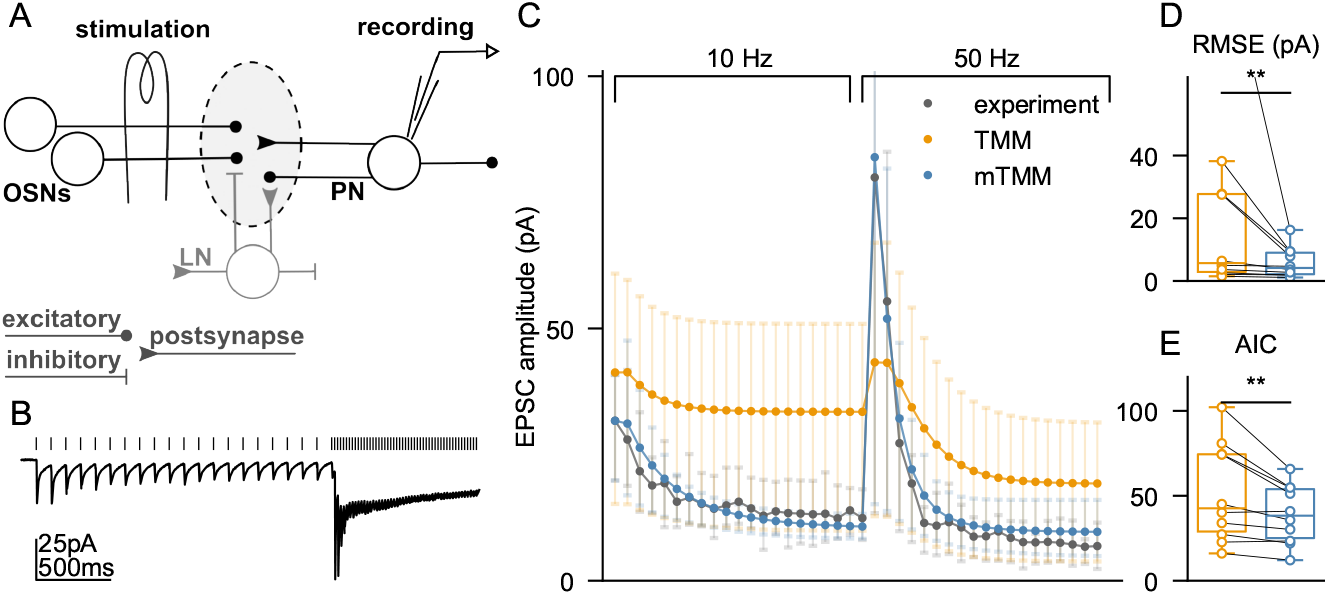
Models of short-term plasticity evaluated at the OSN-PN wild-type synapse. (A) Schematic of the experimental setup. Electrical stimulation of the antennal nerve activates OSN axons, postsynaptic EPSCs are monitored in single PNs using whole-cell patch clamp recording. Adapted from (32). (B) Top: The stimulation protocol assembles a 10 Hz pulse train (100ms inter-stimulus interval), immediately followed by a 50 Hz pulse train (20ms inter-stimulus interval). Bottom: Mean postsynaptic current trace in PNs. Adapted from (25). (C) Analysis of successive EPSC amplitudes (mean *±*25th and 75th percentiles across 10 PNs) in response to the stimulus protocol for experimental recordings (black), original TMM (orange), and the modified TMM (blue). (D) Root Mean Squared Error (RMSE) of the model fit to experimental EPSC amplitudes for TMM and mTMM. (E) Akaike Information Criterion (AIC) comparing TMM with mTMM.

**Table 1.** Optimized model parameters for the TMM and the mTMM.

|  | TMM |  |  |  | mTMM |  |  |
| --- | --- | --- | --- | --- | --- | --- | --- |
|  | min | max | optimal |  | min | max | optimal |
| **A** | 2 | 1024 | 330±295 | **X** | 0.01 | 0.99 | 0.37±0.12 |
| **U** | 0.01 | 0.99 | 0.17±0.13 | **U** | 0.01 | 0.99 | 0.31±0.29 |
| **tau.f** | 12 | 2380 | 46.5±16.2 | **tau.f** | 12 | 2380 | 15.8±4.7 |
| **tau.d** | 12 | 2380 | 510±323 | **tau.d** | 12 | 2380 | 1490±653 |

Model parameters were optimized using an exponential grid search with 10 values per parameter to identify the best-performing parameter sets. Optimal values (mean±standard deviation across 10 PNs) correspond to the parameter set that yielded the lowest weighted error relative to the experimental data. The time constants tau.f and tau.d are reported in ms.

### Modifications improve model performance

Out of this reason, we introduced two modifications to the original model. In the modified TMM (mTMM), the amount of decrease in *x* is defined by (*u*^−^)*X*. The introduction of the exponent *X <* 1 implies a convex dependence of depression on the release probability *u*, meaning that the amount of depression increases with the availability of ready-releasable vesicles in a sublinear manner and approaches a plateau as release probability nears its maximum. Additionally, *u* increases from its initial value *u*_0_ *>* 0 and is incremented immediately after the occurrence of an AP, allowing for the decoupling of the baseline release probability from the activity-dependent increase in release probability (33, 34). The EPSC amplitude is then defined as

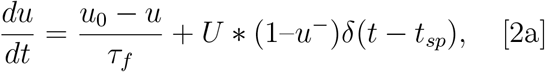

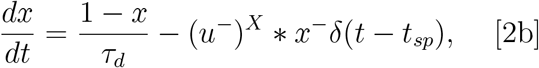

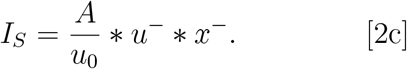

Importantly, the proposed model adjustments in the tuned mTMM allowed to accurately reproduce EPSC amplitudes throughout the stimulation protocol (Tab. 1, Fig. 1 C). The RMSE is strongly and significantly decreased in the mTMM compared to the original model (Fig. 1 D). The Akaike Information Criterion (AIC) is also reduced (Fig. 1 E), indicating that the performance gain by the mTMM is not caused by overfitting. We conclude that the mTMM outperforms the original TMM.

### Model investigation of Unc13A and Unc13B isoforms

In the previous study by Fulterer and colleagues (25), the roles of Unc13A and Unc13B isoforms in synaptic transmission at the OSN–PN synapse were investigated in separate knock-down (KD) experiments of both isoforms (Fig. 2). We re-analyzed these data with the objective to reveal the change of model parameters when one isoform predominates. For parameter optimization we employed a Bayesian method that resulted in a four-dimensional likelihood distribution across the four adjustable model parameters (33). We then used the parameter set with the highest likelihood for the optimal model, which accurately replicated the experimentally observed EPSC amplitudes in all three experimental conditions (Fig. 2 B,E,H).

**Fig. 2.**
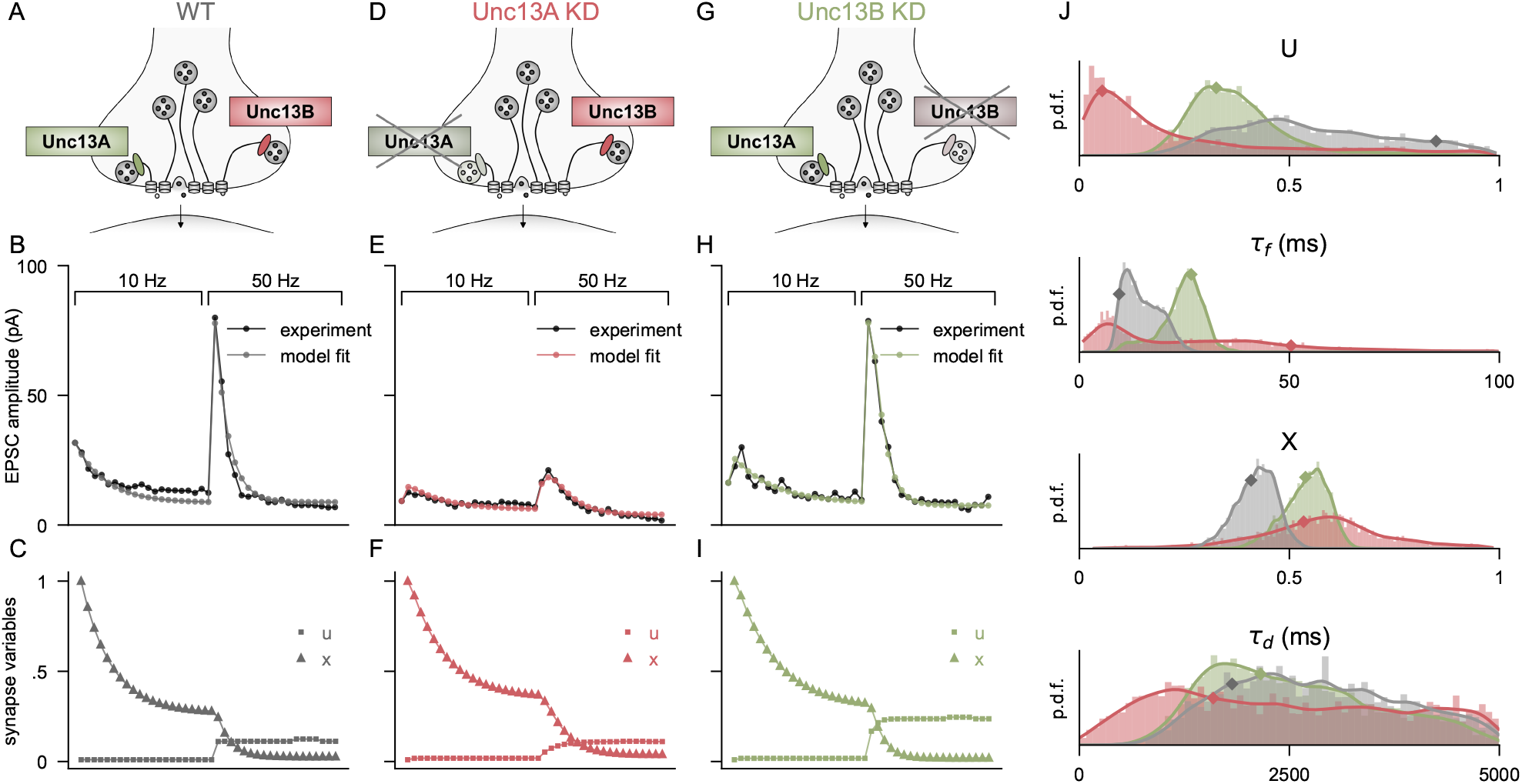
Tuning the extended mTMM to Unc13 KD conditions. (A) Schematic of vesicle release mediated by the Unc13A and Unc13B isoforms. Model fit of the extended mTMM to the mean EPSC amplitudes under wild-type (WT) conditions. Experimental data adapted from (25). (C) Dynamics of short-term plasticity variables x and u in the WT condition. (D–F) Corresponding panels for the Unc13A KD condition, showing (D) vesicle release via Unc13A, (E) model fit to EPSC amplitudes, and (F) short-term plasticity dynamics. (G–I) As in panels D–F, but for the Unc13B KD condition. (J) Histograms and kernel density estimates of posterior distributions of synaptic parameters for WT (grey), Unc13A KD (red), and Unc13B KD (green). Diamonds indicate the maximum *a posteriori* estimates, corresponding to the highest log-likelihood values that were used for fits in B,E,H.

Data and model clearly show a strong effect of Unc13A KD in comparison to the WT synapse and only a weak effect of the Unc13B KD, confirming the result that the OSN-PN synapse is Unc13A dominated. The Unc13A KD is expressed in a strong reduction of the paired-pulse facilitation at the transition from 10Hz to 50Hz stimulation. In our model, this finds expression in a strong reduction of the parameter *U* in Fig. 2J, while the marginal distribution for the facilitation time constant *τ*_*f*_ is now flattened. The depression time constant *τ*_*d*_ assumes smaller values, indicating a faster recovery of *x* and thus clear depression takes effect only for short inter-pulse intervals. Together, the model parameters allow us to infer that Unc13A supports the phasic and Unc13B the tonic component of synaptic transmission.

## Discussion

This study presents a tailored model of STP that captures the functional diversity introduced by Unc13 isoforms at central synapses in the fly olfactory system.

### Functional implications

Previous studies have shown that the distinct roles of Unc13 isoforms in shaping synaptic transmission have important functional consequences across multiple levels of neural processing, as evidenced in *D. melanogaster* (22). These isoform-specific differences extend beyond synaptic physiology to influence sensory processing and behavior. For instance, Unc13A and Unc13B differentially shape sensory coding and innate behavior towards appetitive and aversive cues, including food odors (35–37). In the context of learned behavior, Unc13A plays a particularly critical role in the formation of shortand mid-term aversive memories (38– 40). Our phenomenological model of Unc13A or Unc13B-dominated synaptic transmission can serve as a modular component for integration into existing circuit models of the fly olfactory system. This offers a framework for investigating the functional implications of molecular properties of synaptic release.

### Comparison to other models

Several previous studies have proposed modified versions of the widely applied TMM, tailored to specific use cases (18, 33, 34). For instance, (34) introduced a version of the TMM in which the equilibrium release fraction *u*_0_ is decoupled from the spike-triggered increment *U*. This modification allows independent tuning of both model parameters, similar to the approach used in this study. In contrast to the phenomenological TMM, other models of synaptic transmission have taken a more mechanistic perspective. For example, several studies have explored parallel or sequential vesicle priming mechanisms, incorporating distinct vesicle states and transitions between loosely and tightly docked vesicles, as well as their replenishment from a reserve pool (41–45). These distinct vesicle pools align with the separate roles of Unc13 isoforms, offering a potential alternative framework for modeling the dual release mechanisms. Another study has highlighted how vesicleCa2+ channel distances shape STP through activity-dependent modulation of the readily releasable pool (46). In this work, we present a phenomenological model capturing Unc13specific diversity in short-term synaptic dynamics. The key advantage of this model lies in its small parameter set, providing a biologically realistic yet computationally efficient mathematical abstraction of the underlying molecular mechanism.

### Limitations

While our proposed model effectively captures the transmission dynamics of the studied synapse type and offers insight into Unc13 isoform function, certain limitations should be considered. First, our study focuses on synapses between OSNs and PNs in the fly olfactory pathway, which are dominated by the phasic Unc13A isoform (Fig. 2,(25)). This allows us to infer properties of Unc13Aspecific transmission but does not directly inform us on the transmission properties of the tonic Unc13B isoform. Future work could explore whether our model captures well Unc13B-dominated transmission, such as at the synapse between local interneurons (LNs) in the fly antennal lobe or between PNs and specific classes of lateral horn neurons (LHNs) (25, 36). While (36) provide initial electrophysiological recordings from PN-LHN synapses, detailed transmission data for Unc13B-rich sites remain limited, highlighting an important area for future experimental and modeling efforts. Second, the experimental data used to fit our model, particularly in the Unc13 KD conditions (Fig. 2), is limited in sample size. Given this constraint and the variability within the dataset and across individuals, we chose to fit our model to the mean EPSC amplitudes rather than individual responses. This approach may overlook sample-specific variations and does not easily allow for statistical analyses between conditions (47). Finally, our model considers synaptic dynamics in a highly simplified and isolated context. For example, we do not incorporate lateral and feedback inhibition from LNs in the antennal lobe that targets OSNs and PNs (48, 49) and may thus influence transmission dynamics.

### Outlook

Future studies can build on our framework by i) comparing the examined Unc13A dominated synapse to differently Unc13B tuned synapse types (25, 36), and ii) exploring how the distinct roles of Unc13 isoforms in shaping synaptic transmission extend to sensory processing and behavior in largescale neural network simulations, highlighting the documented functional significance of Unc13 isoform-specific transmission dynamics in *D. melanogaster* (22).

## Materials and Methods

### Experimental data

The experimental data were obtained from (25). A single *in vivo* recording was performed per animal. In each PN, EPSC peak amplitudes were extracted for all stimulus pulses.

### Model

Our minimal spiking neural network was implemented using Brian2. The model included a SpikeGeneratorGroup(), which mimicked the stimulation protocol from the experimental dataset, and a synapse with either the original or modified variant of the TMM connected to a postsynaptic neuron.

### Parameter optimization

To optimize the model, we conducted a grid search to determine for which parameter set the TMM and mTMM could capture the experimental synaptic dynamics for each single recording optimally. For parameters *X* and *U*, we selected a uniformly spaced grid, whereas for *A, τ*_*f*_, and *τ*_*d*_ we used an exponential grid to efficiently sample a broader range, each with a grid size of 10. For each parameter set we calculated the root mean square error (RMSE) between simulated EPSC amplitudes *y*_*i*_ and experimental EPSC amplitudes *ŷ*_*i*_ across 40 stimulations as

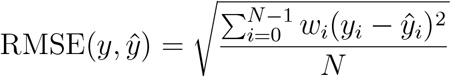

where a weighted error was used, with weights *w*_*i*_ proportional to the experimental peak value in order to proportionally emphasize high peak values observed after the frequency change from 10 to 50 Hz.

To ensure comparability of the model performance for the TMM and the mTMM considering the fact that the latter has one additional parameter, we applied the Akaike Information Criterion (AIC)

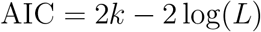

where *L* is the model likelihood and *k* is the number of free model parameters. Lower AIC values indicate a model that better fits the data while penalizing excess complexity.

### Bayesian inference

For parameter tuning and comparison of the KD conditions, we used Bayesian optimization (33). Due to the limited sample sizes in the KD experiments (n=7 in Unc13A KD, n=5 in Unc13B KD), we used the mean postsynaptic EPSC amplitudes. We assumed uniform prior distributions for the parameters *X, U, τ*_*f*_, *τ*_*d*_. We assumed Gaussian noise, with mean *µ* equal to the average amplitude per stimulus pulse. Due to limited sample size, the standard deviation was estimated visually, leading to a selection of *σ* = 10 *pA*. Sampling employed the Differential Evolution Metropolis variant, DEMetropolisZ (50), with a burn-in of 2500 samples, followed by 7500 samples across four chains. The optimal parameter set for each condition was selected based on the maximum log-likelihood.

All data analysis and parameter fitting were performed in Python 3. The programming code is accessible at: https://github.com/nawrotlab/Unc13-STP-Model.

## ACKNOWLEDGMENTS

We thank Katherine Eyring and Katherine Nagel for providing us with the electrophysiological data set as analyzed in (25). Funding was received from the German Research Foundation (DFG) through the priority program “Evolutionary Optimization of Neuronal Processing” (DFG-SPP 2205, ID 430592330 to M.P.N.), the Collaborative Research Center “Motor Control in Health and Disease” (DFG-SFB 1451, Project A06, ID 431549029 to M.P.N.), and the Research Unit “Structure, Plasticity and Behavioral Function of the Drosophila Mushroom Body” (DFG-FOR 2705, ID 365082554 to S.S. and M.P.N.).

## Notes

### Competing Interest Statement

The authors have declared no competing interest.

